# Genotyping by sequencing for identification and mapping of QTLs for bioenergy-related traits in sweet sorghum

**DOI:** 10.1101/2020.06.25.170662

**Authors:** Kanokwan Teingtham, David M. Braun, Ismail Dweikat

**Affiliations:** Department of Agronomy and Horticulture, University of Nebraska-Lincoln, 202 Keim Hall, Lincoln, NE 68583, USA; Division of Biological Sciences, University of Missouri, 110 Tucker Hall, Columbia, MO 65211, USA

**Keywords:** *Sorghum bicolor* (L.) Moench, Recombinant inbred lines, ICiMapping, Sugar transporter gene

## Abstract

Sweet sorghum (*Sorghum bicolor* L. Moench) is a promising bioenergy crop. To increase the productivity of this crop, marker-assisted breeding will be important to advance genetic improvement of sweet sorghum. The objective of the present study was to identify quantitative trait loci (QTLs) associated with bioenergy-related traits in sweet sorghum. We used 188 F_7_ recombinant inbred lines (RILs) derived from a cross between sweet sorghum (Wray) and grain sorghum (Macia). The RILs and their parental lines were grown at two locations in 2012 and 2013. Genotyping-by-sequencing analysis of the RILs allowed the construction of a map with 979 single nucleotide polymorphisms. Using the inclusive composite interval mapping of additive QTLs, major QTLs for flowering time and head moisture content were detected on chromosome 6, and explained 29.45% and 20.65% of the phenotypic variances (PVE), respectively. Major QTLs for plant height (29.51% PVE) and total biomass yield (16.46% PVE) were detected on chromosome 7, and QTLs for stem diameter (9.43% PVE) and 100 seed weight (22.97% PVE) were detected on chromosome 1. A major QTL for brix (39.92% PVE) and grain yield (49.14%) PVE co-localized on chromosome 3, was detected consistently across four environments, and is closely associated with a SWEET sugar transporter gene. Additionally, several other QTLs for brix identified in this study or reported previously were found to be associated with sugar transporter genes. The identified QTLs in this study will help to further understand the underlying genes associated with bioenergy-related traits and could be used for development of molecular markers for marker-assisted selection.

## Introduction

Bioenergy is an alternative and renewable energy derived from biomass resources, which includes waste (agricultural production waste, mill wood waste, urban organic waste, etc.), standing forests, and agricultural crops (Demirbaş 2001). It is expected that bioenergy will provide 30% of the world’s energy by 2050 (Guo et al. 2015). Sweet sorghum (*Sorghum bicolor* L. Moench) is a leading bioenergy crop (Calviño and Messing 2012; Carpita and McCann 2008; Zegada-Lizarazu and Monti 2012). Sweet sorghum is highly productive, with low input requirements, and is drought-tolerant (Rooney et al. 2007; Zegada-Lizarazu and Monti 2012). Sweet sorghum accumulates large amounts of carbohydrates in its stalk and produces total biomass as high as 30 Mg ha^-1^ (Bihmidine et al. 2015; Rooney et al. 2007). Stalk carbohydrates are easily converted to ethanol via fermentation of stalk juice. The pressed stalk, after juice extraction, can be compressed into pellets, which are combustible. Thus, sweet sorghum could be used for both biofuel and thermo-electrical energy production (Zegada-Lizarazu and Monti 2012).

To be an efficient energy crop, sweet sorghum should be genetically improved. Biotechnology, genomics, and marker-assisted breeding will be important for genetic improvement of sweet sorghum (Madhusudhana 2014; Rooney et al. 2007). Genomic regions linked to complex traits can be identified by genetic mapping and QTL (quantitative trait locus) analysis (Shehzad and Okuno 2014). The fundamental idea underlying QTL analysis is to associate genotype and phenotype in a population exhibiting a genetic variation (Broman and Sen 2009). Analysis of QTLs is the first strategy in marker-assisted selection (MAS) for phenotypic traits (Shehzad and Okuno 2014).

QTLs for bioenergy-related traits in sweet sorghum have been reported, including flowering time (Felderhoff et al. 2012; Murray et al. 2008a; Murray et al. 2008b; Ritter et al. 2008; Shiringani et al. 2010), plant height (Felderhoff et al. 2012; Guan et al. 2011; Murray et al. 2008a; Murray et al. 2008b; Ritter et al. 2008; Shiringani et al. 2010), biomass (Felderhoff et al. 2012; Murray et al. 2008a; Shiringani and Friedt 2011, stem diameter (Shiringani et al. 2010), sugar content (Bian et al. 2006; Felderhoff et al. 2012; Guan et al. 2011; Lv et al. 2013; Murray et al. 2008a; Murray et al. 2008b; Ritter et al. 2008; Shiringani et al. 2010), grain yield (Felderhoff et al. 2012; Murray et al. 2008b; Ritter et al. 2008), and seed weight (Murray et al. 2008b). However, many genes influencing biomass productivity of sweet sorghum remain unknown. Recently, the application of next-generation sequencing technologies has led to whole genome sequencing, making Genotype-by-sequencing (GBS) feasible for various species (Elshire et al. 2011). GBS has been used to generate single-nucleotide polymorphisms (SNPs) and QTL mapping (Beissinger et al. 2013). Using GBS for QTL mapping improves sensitivity and resolution of QTL detection (Kong et al. 2018; Verma et al. 2015).

Thus, the objectives of this study were 1) to identify QTLs associated with bioenergy-related traits in sweet sorghum using GBS, and 2) to confirm previously identified QTLs in an independent genetic background. QTLs for the bioenergy-related traits of sweet sorghum: flowering time, plant height, head moisture content, biomass yield, stem diameter, brix (sugar content), grain yield, and 100 seed weight were detected for 188 F_7_ recombinant inbred lines (RILs) derived from a cross between Macia (grain sorghum) and Wray (sweet sorghum) (Bihmidine et al. 2015).

These QTLs could help identify genes that influence biomass productivity of sweet sorghum, and could be used for development of molecular markers to select individuals with valuable bioenergy-related traits within a breeding population using these markers.

## Materials and Methods

### Plant material

Two hundred F_6_ and F_7_ RILs derived from a cross between Macia and Wray were used for the QTL evaluation in this study. Macia (SDS 3220, PI 565121) is a grain sorghum developed by the Botswana National Agricultural Research System (Saadan et al. 2000). Macia is 1.4-1.6 m tall, with 7.3% stalk sugar content, grain yield about 3.3 Mg ha^-1^, and biomass yield about 10.1 Mg ha^-1^ (Makanda et al. 2009; Setimela et al. 1997). Wray is a sweet sorghum developed by U.S. sugar-breeding programs (Broadhead et al. 1978; Hills et al. 1990). Wray (PI 653616) is 2.33 m tall, with 19% stalk sugar content, grain yield about 1.1 Mg ha^-1^, and biomass yield about 22.2 Mg ha^-1^ (Bihmidine et al. 2015; Pedersen et al. 2013).

### Field experiment

A total of 210 lines, including 200 RILs, two parental lines, and eight check cultivars (RTx430 (PI 655996), BTx623 (NSL 466845), San Chi San (PI 542718), N586 (PI 642391), Simon, UNL 3016, UNL 4016, and UNL 26297B) were planted under rainfed conditions at University of Nebraska-Lincoln experimental farms at Havelock and Mead, Nebraska in the summer seasons of 2012 and 2013. In 2012, the planting dates were May 23^rd^ at Havelock and May 25^th^ at Mead. In 2013, the planting dates were May 24^th^ at Havelock and May 23^rd^ at Mead. The experimental design was an alpha lattice incomplete block design with 15 incomplete blocks of 14 plots each per replication (15×14 alpha lattice), with two replications in each environment. The total was 420 plots for each environment. A plot was a single row measuring 4.5 m long, with 0.75 m between rows. A single row was sown at a rate of 50 seeds per row.

### Phenotyping of bioenergy-related traits

1) Flowering time was measured as the duration of days from planting until 50% of the plants within a plot were shedding pollen. In 2012, harvesting dates were September 18^th^ - 26^th^ at Havelock and October 2^nd^ - 5^th^ at Mead. In 2013, harvesting dates were September 23^rd^ - 27^th^ at Havelock and October 1^st^ - 9^th^ at Mead. On average, harvesting was performed 125 days after planting. At harvesting times, five middle plants within a plot were arbitrarily selected to measure plant height, total biomass, stem diameter, brix (sugar content in stem juice), head and stem moisture content, grain yield, and 100 seed weight. 2) Plant height was measured as the distance in cm from the base of the plant to the tip of the panicle. 3) Total biomass in Mg ha^-1^ was collected when plants reached their physiological maturity by cutting the plants near the soil surface and separating them into panicles (heads), leaves, bottom stems (the length of 0-20 cm above ground), and remaining stems. The sub-samples were bagged and weighed immediately to obtain the wet weight and placed into an oven at 120-160°C for ten days to completely dry the samples. Dried subsamples were reweighed and were summed up to obtain the total dry weight. Total biomass was calculated as follows:

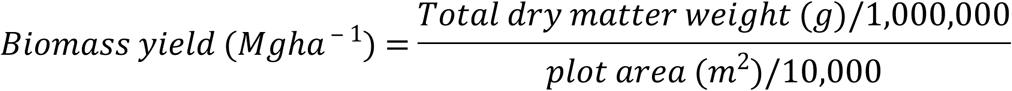

4) Stem diameter (cm), and 5) brix readings were obtained from the bottom stem sections before drying. Brix degree (°Brix) as a measure of soluble solids (mainly sucrose) in the stem juice was measured using a hand-held refractometer (MASTER-T Brix 0-32% with ATC, Atago Co., LTD, Tokyo, Japan). Because of the high volume workload in the field, stem diameter and brix readings were measured in the lab within 2-3 days. The bottom stems were kept in covered plastic boxes and put in a cold room (4°C) to minimize sugar metabolism and to maintain brix level until the measurement. 6) Head moisture content and 7) stem moisture content were calculated as the percentage difference between wet biomass and dry weight. 8) Grain yield was calculated as follows:

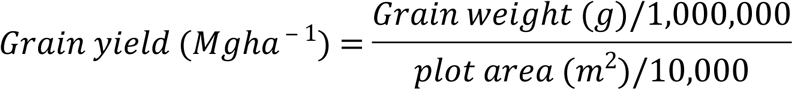

Sub-samples of panicles were threshed and cleaned. Seeds were randomly sampled for 9) 100 seed weight (g).

### Phenotypic Analysis

Analysis of variance (ANOVA) for bioenergy-related traits was performed for each environment and combined environments using the PROC MIXED procedure (Littell et al. 2006) of SAS version 9.4 (SAS Institute, 2008). Pearson’s correlation coefficients between traits were calculated using the PROC CORR procedure of SAS. Frequency distribution for bioenergy-related traits was performed for each environment using the PROC CAPABILITY. Narrow-sense heritability with the standard error was determined using SAS code which is available at http://www4.ncsu.edu/~jholland/heritability/Inbreds.html (Holland et al. 2003).

### Genotyping-by-sequencing (GBS)

A total of 190 genomic DNA samples, including 188 randomly selected RILs from 200 RILs and two parental lines (Macia and Wray), were sent to the Institute for Genomic Diversity, Cornell University for GBS. The DNA samples were genotyped by GBS (Elshire et al. 2011) using the Illumina HiSeq 2000/2500 (100 bp, single-end reads). The 190 genomic DNA samples were isolated from 20-day-old leaf tissue using a magnetic bead DNA extraction method (Xin and Chen 2012). The DNA samples were diluted to 100 ng/µl and 40 µl of each sample submitted for GBS. The raw sequence reads (FASTQ files) were analyzed using the TASSEL4 GBS pipeline (Bradbury et al. 2007). The raw sequence reads were aligned to the reference sequence for sorghum (Sbi1) (Paterson et al. 2009) using Burrows-Wheeler Aligner, BWA (Li and Durbin 2009). The formatted reference sequence file for BWA was provided by Illumina. The TASSEL4 GBS pipeline provided the HapMap file containing high-throughput SNPs (10,424 SNPs). The HapMap file was imported into Microsoft Excel 2010 for coding markers following ICiMapping format (Wang et al. 2012).

### Linkage map construction

Linkage maps were constructed by IciMapping 3.2 (Wang et al. 2012). Before linkage map construction, binning of redundant markers, using the ICiMapping BIN function was performed to calculate the missing rate, and linkage maps were pre-constructed to evaluate a 1:1 segregation ratio of markers using the chi-square test (P≤0.01). SNPs with ≥80% missing rate and significant SNPs from the chi-square test were removed from the data set. Finally, genetic linkage maps were reconstructed with 979 selected SNPs.

### QTL analysis

QTLs analysis, on biparental populations (BIP), was performed by IciMapping 3.2. Inclusive composite interval mapping of additive QTLs (ICIM-ADD) was used to conduct QTL mapping. Genetic linkage maps, means of phenotypic data for bioenergy-related traits for 188 RILs across four environments, and genotypic data of 979 SNPs were used for QTL analysis. The Kosambi mapping function was used to convert recombination frequency to mapping distance. Step in scanning was assigned at one centiMorgan (cM). A thousand-permutation test was applied to each data set to determine LOD (log_10_ of the likelihood odds ratio) threshold (P≤0.01), with LOD score >3 was used to determine significant QTLs. Phenotypic variation explained by the marker (%PVE) and the additive effect of QTLs were estimated by IciMapping.

## Results

### Phenotypic data

Significant differences (P<0.05) were observed among genotypes and genotype x environment interactions for all traits measured in each environment and combined environments (Table 1). Wray had greater means than Macia for all traits except stem diameter, grain yield, and 100 seed weight. The mean values for all traits of RILs were between the parental lines values (Table 1; Fig.1). The narrow-sense heritability (*h*^*2*^), estimated for all traits and environments, ranged from 0.14 for stem moisture content at Havelock in 2013 to 0.96 for plant height at Havelock in 2013 (Table 1).

**Table 1.**
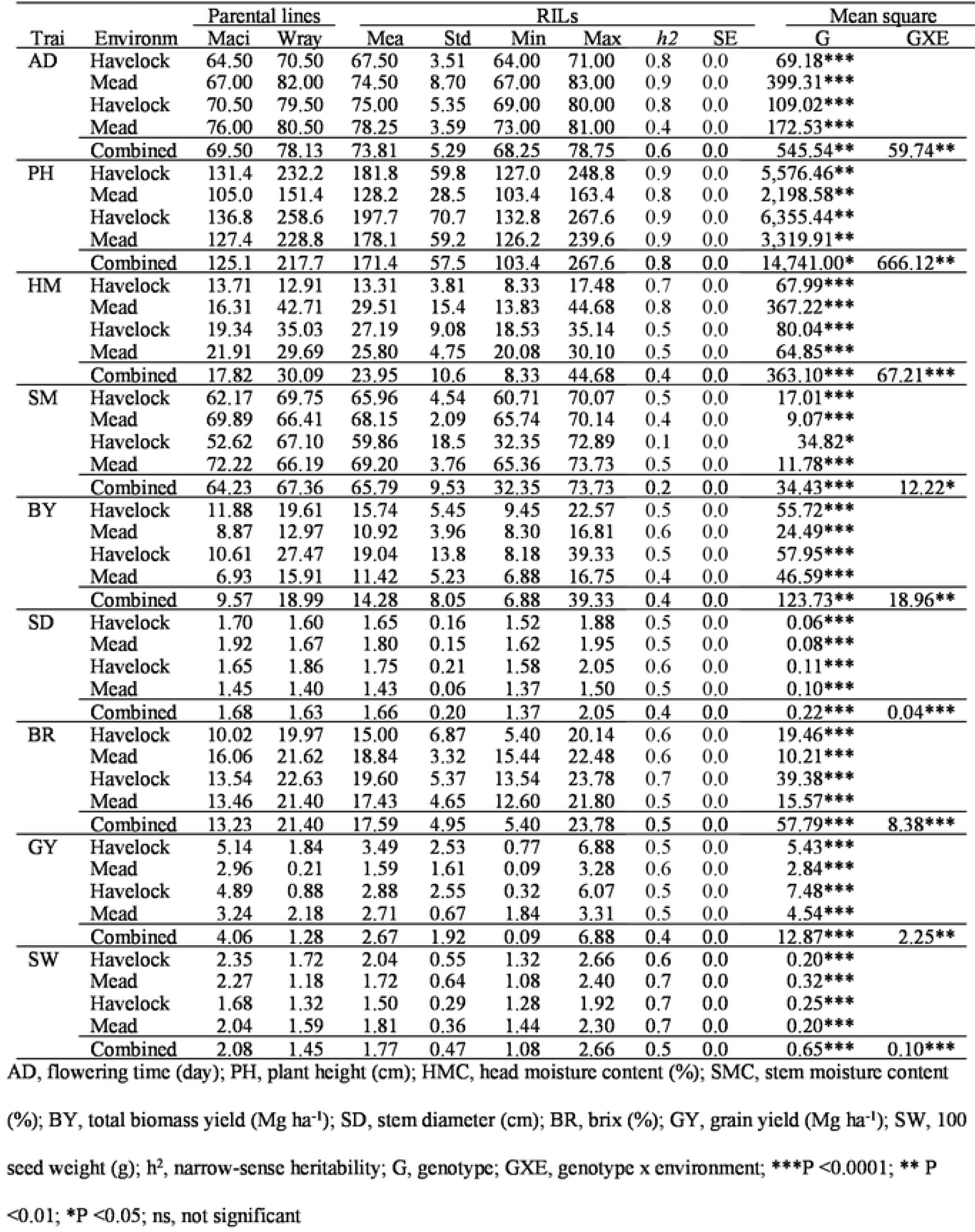
Descriptive statistics, ANOVA, and heritability estimates of nine trails of parental lines and RILs across four environments

**Fig. 1.**
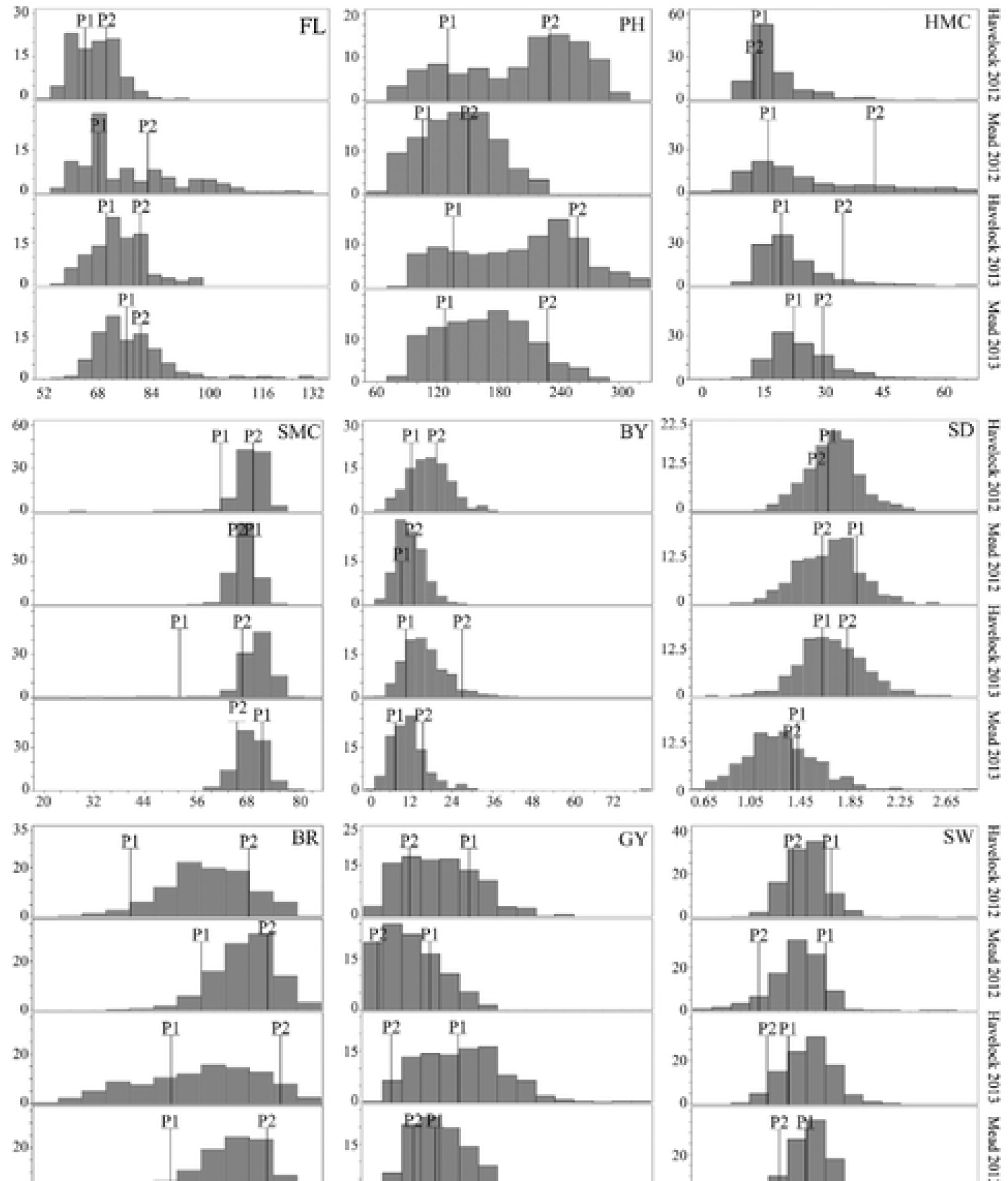
Frequency distribution of nine bioenergy-related traits of RILs for four environments. Pl, parental line 1 (Macia); P2, parental line 2 (Wray); FL, flowering time (day); PH, plant height (cm); HMC, head moisture content (%); SMC, stem moisture content (%); BY, tota1biomass yield (Mg ha^-1^); SD, stem diameter(cm); BR, brix (%); GY, grain yield (Mg ha^-1^); SW, 100 seed weight (g)

The highest positive correlation was observed between flowering time and head moisture content (*r*=0.73) (Table 2). Sweet sorghum lines with later flowering time had higher head moisture content. Total biomass yield was positively correlated with all traits except head moisture content (Table 2). Total biomass yield correlated best with plant height (*r*=0.67). Brix was positively correlated with head moisture content (*r*=0.06) and 100 seed weight (*r*=0.15) and was negatively correlated with stem moisture content (*r*=-0.41), grain yield (*r*=-0.39), and stem diameter (*r*=-0.13) (Table 2).

**Table 2.**
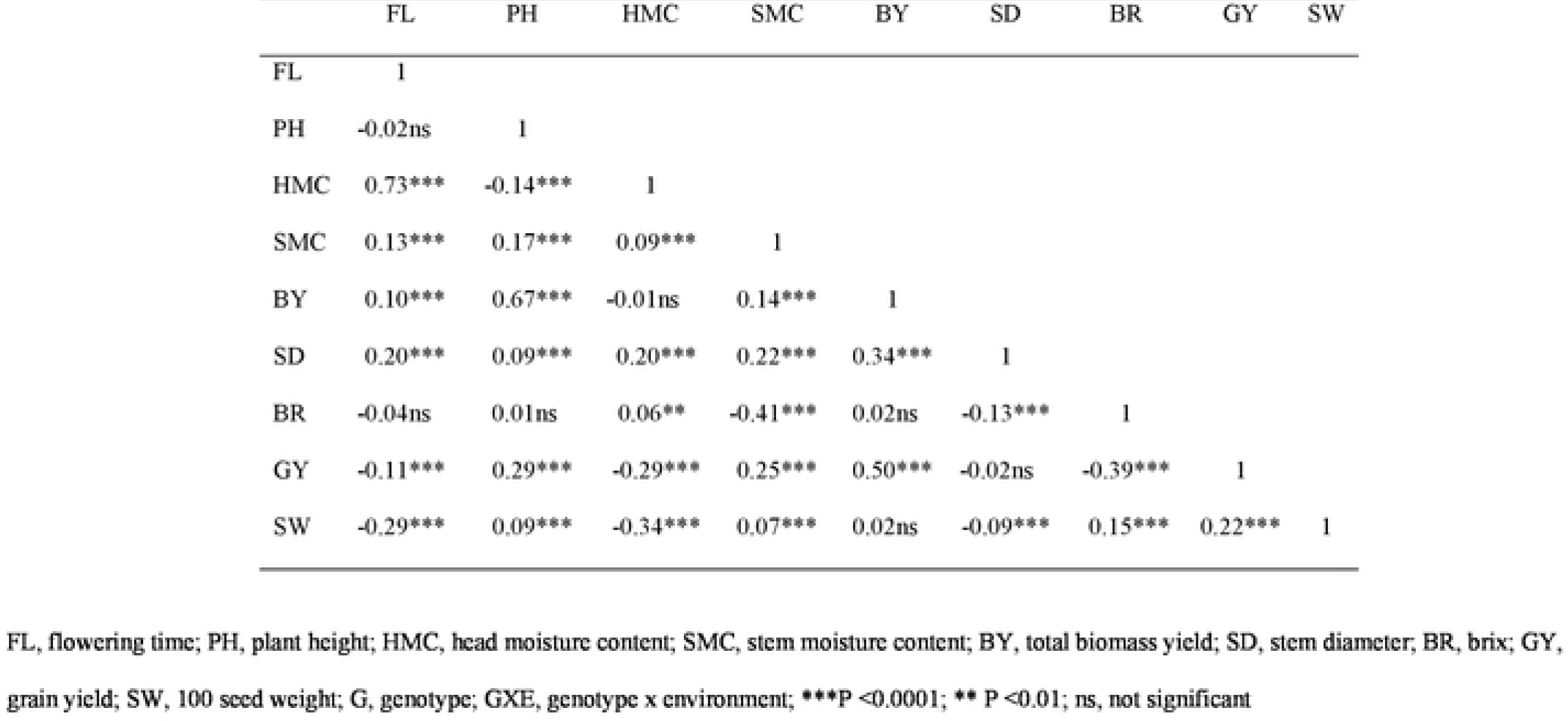
Correlation coefficients among traits based on 1east square means in parental lines and RILs across four environments

|  | FL | PH | HMC | SMC | BY | SD | BR | GY | SW |
| --- | --- | --- | --- | --- | --- | --- | --- | --- | --- |
| FL | 1 |  |  |  |  |  |  |  |  |
| PH | -0.02ns | 1 |  |  |  |  |  |  |  |
| HMC | 0.73*** | -0.14*** | 1 |  |  |  |  |  |  |
| SMC | 0.13*** | 0.17*** | 0.09*** | 1 |  |  |  |  |  |
| BY | 0.10*** | 0.67*** | -0.01ns | 0.14*** | 1 |  |  |  |  |
| SD | 0.20*** | 0.09*** | 0.20*** | 0.22*** | 0.34*** | 1 |  |  |  |
| BR | -0.04ns | 0.01ns | 0.06** | -0.41*** | 0.02ns | -0.13*** | 1 |  |  |
| GY | -0.11*** | 0.29*** | -0.29*** | 0.25*** | 0.50*** | -0.02ns | -0.39*** | 1 |  |
| SW | -0.29*** | 0.09*** | -0.34*** | 0.07*** | 0.02ns | -0.09*** | 0.15*** | 0.22*** | 1 |

FL, flowering time; PH, plant height; HMC, head moisture content; SMC, stem moisture content; BY, total biomass yield; SD, stem diameter; BR, brix; GY, grain yield; SW, 100 seed weight; G, genotype; GXE, genotype x environment; \*\*\*P <0.0001; \*\* P <0.01; ns, not significant

### Genotyping-by-sequencing (GBS)

Sequencing yielded approximately 1.66 million reads in total for the 190 genotypes (Table 3). About 1.32 million reads (79.76%) were successfully mapped to the sorghum reference genome (Sbi1) (Table 3). The unmapped reads included reads that aligned to multiple positions (0.20 million reads, 12.06%) and reads that could not be aligned (0.14 million reads, 8.18%) (Table 3).

**Table 3.** Alignment results of DNA tags (reads) of l90 genomic DNA samples

| Tags on Physical Map | Number of tags | % |
| --- | --- | --- |
| Tags were aligned to unique positions | 1,322,720 | 79.76 |
| Tags were aligned to multiple positions | 199,995 | 12.06 |
| Tags could not be aligned | 135,704 | 8.18 |
| Total | 1,658,419 | 100.00 |

### Linkage map construction

A total of 979 SNPs from GBS were mapped onto 10 linkage groups (chromosomes) based on the physical position of the SNPs. The linkage map spanned a total of 1,707.11 cM with an average inter-marker distance 1.74 cM (Table 4). The maximum distance between markers was 47.11 cM (Chr 8) (Table 4). On average, each linkage group contained 98 markers that spanned an average of 170.71 cM. The length of the linkage groups ranged from 87.63 cM (Chr 9) to 212.03 cM (Chr 2) (Table 4).

**Table 4.** Description of basic characteristics of a genetic linkage map of Macia x Wray population using SNPs generated from genotyping-by-sequencing (GBS)

| Chr | No. of Markers | Distance between markers (cM) |  |  |  |
| --- | --- | --- | --- | --- | --- |
|  |  | Total | Min | Max | Average |
| 1 | 109 | 199.16 | 0 | 10.35 | 1.83 |
| 2 | 164 | 212.03 | 0 | 8.99 | 1.29 |
| 3 | 129 | 210.47 | 0 | 17.79 | 1.63 |
| 4 | 55 | 138.21 | 0 | 16.08 | 2.51 |
| 5 | 97 | 208.83 | 0 | 32.49 | 2.15 |
| 6 | 88 | 168.56 | 0 | 42.10 | 1.92 |
| 7 | 87 | 127.75 | 0 | 26.20 | 1.47 |
| 8 | 71 | 219.82 | 0 | 47.11 | 3.10 |
| 9 | 85 | 87.63 | 0 | 9.97 | 1.03 |
| 10 | 94 | 134.65 | 0 | 9.84 | 1.43 |
| Total | 979 | 1,707.11 | - | - | - |
| Average | 97.9 | 170.71 | 0 | 22.09 | 1.74 |

### QTL analysis

QTLs for nine bioenergy-related traits were analyzed, including flowering time, plant height, head moisture content, stem moisture content, biomass yield, stem diameter, brix, grain yield, and 100 seed weight. A total of 29 QTLs for eight of nine traits were detected for combined environments using ICIM-ADD. None of the QTLs for stem moisture content were detected for combined environments. The QTL locations for four environments and combined environments are presented in Table 5, Fig. 2, and Fig. 3. The QTLs detected for combined environments were dispersed across chromosomes 1, 2, 3, 4, 6, and 7. The QTLs explained 3.10-49.14% of the PVE, with a LOD range of 4.28-30.46, and an interval of 0.33-17.79 cM.

**Table 5.** QTIs identified for eight traits in Macia x Wray RIL mapping population for combined environments

| Trait | QTL | Chr | Left Marker<br>Marker | Right Marker<br>Marker | Left<br>Position | Right<br>Position | Peak<br>Position | Interval<br>(cM) | LOD | PVE<br>(%) | Add |
| --- | --- | --- | --- | --- | --- | --- | --- | --- | --- | --- | --- |
| Flowering time | qFL1 | 1 | S1_19760428 | S1_20278787 | 59.63 | 62.88 | 61 | 3.25 | 13.18 | 18.17 | 3.63 |
|  | qFL6 | 6 | S6_5982269 | S6_41973878 | 0 | 6.89 | 4 | 6.89 | 18.62 | 29.45 | -4.66 |
| Plant height | qPH1 | 1 | S1_10103911 | S1_10116667 | 6.94 | 7.27 | 7 | 0.33 | 4.88 | 3.10 | 8.04 |
|  | qPH6 | 6 | S6_41974107 | S6_42311647 | 6.89 | 8.56 | 8 | 1.67 | 18.18 | 14.42 | -17.53 |
|  | qPH7_1 | 7 | S7_54778890 | S7_55156277 | 56.16 | 57.53 | 57 | 1.37 | 30.46 | 29.51 | -24.84 |
|  | qPH7_2 | 7 | S7_58245093 | S7_58601476 | 82.13 | 87.37 | 86 | 5.24 | 24.48 | 22.34 | -21.68 |
| Head moisture<br>content | qHMC1_1 | 1 | S1_15037202 | S1_15394276 | 32.81 | 34.92 | 34 | 2.11 | 4.67 | 7.97 | 1.81 |
|  | qHMC1_2 | 1 | S1_20321046 | S1_24666533 | 63.83 | 71.38 | 65 | 7.55 | 4.44 | 7.15 | 1.73 |
|  | qHMC6 | 6 | S6_41974107 | S6_42311647 | 6.89 | 8.56 | 7 | 1.67 | 12.95 | 20.65 | -2.92 |
| Biomass yield | qBY4 | 4 | S4_57956001 | S4_58857380 | 105.07 | 107.35 | 106 | 2.28 | 4.40 | 5.72 | -0.97 |
|  | qBY6 | 6 | S6_5982269 | S6_41973878 | 0 | 6.89 | 4 | 6.89 | 8.29 | 10.93 | -1.35 |
|  | qBY7_1 | 7 | S7_54507788 | S7_54744818 | 53.7 | 54.84 | 54 | 1.14 | 10.74 | 13.29 | -1.47 |
|  | qBY7_2 | 7 | S7_58245093 | S7_58601476 | 82.13 | 87.37 | 86 | 5.24 | 12.22 | 16.46 | -1.64 |
| Stem diameter | qSD1_1 | 1 | S1_19423098 | S1_19760428 | 57.36 | 59.63 | 58 | 2.27 | 7.52 | 9.26 | 0.05 |
|  | qSD1_2 | 1 | S1_19760428 | S1_20278787 | 59.63 | 62.88 | 61 | 3.25 | 7.78 | 9.43 | 0.05 |
|  | qSD1_3 | 1 | S1_57417126 | S1_57674671 | 122.07 | 126.40 | 124 | 4.33 | 6.49 | 7.91 | 0.05 |
|  | qSD4 | 4 | S4_5042376 | S4_7171350 | 24.1 | 29.24 | 26 | 5.14 | 6.03 | 7.71 | 0.05 |
|  | qSD6 | 6 | S6_57834516 | S6_57863270 | 148.42 | 149.16 | 149 | 0.74 | 4.38 | 4.81 | -0.04 |
| Brix | qBR2 | 2 | S2_13246714 | S2_14023143 | 56.29 | 59.40 | 59 | 3.11 | 4.28 | 5.28 | -0.62 |
|  | qBR3_1 | 3 | S3_62297204 | S3_68877230 | 158.76 | 176.55 | 169 | 17.79 | 19.14 | 39.92 | -1.71 |
|  | qBR3_2 | 3 | S3_68877230 | S3_69154987 | 176.55 | 178.95 | 177 | 2.40 | 13.48 | 19.52 | -1.20 |
|  | qBR4_1 | 4 | S4_56025428 | S4_57635417 | 94.27 | 100.68 | 97 | 6.41 | 5.84 | 9.05 | 0.82 |
|  | qBR4_2 | 4 | S4_57635417 | S4_57727357 | 100.68 | 101.97 | 101 | 1.29 | 6.52 | 8.45 | 0.79 |
|  | qBR7 | 7 | S7_58601476 | S7_60256664 | 87.37 | 95.91 | 90 | 8.54 | 4.33 | 6.17 | -0.67 |
| Grain yield | qGY3 | 3 | S3_62297204 | S3_68877230 | 158.76 | 176.55 | 170 | 17.79 | 18.99 | 49.14 | 0.97 |
| 100 seed weight | qSW1 | 1 | S1_62441638 | S1_62574013 | 148.7 | 149.25 | 149 | 0.55 | 16.75 | 22.97 | 0.13 |
|  | qSW4 | 4 | S4_57956001 | S4_58857380 | 105.07 | 107.35 | 107 | 2.28 | 7.33 | 8.03 | 0.08 |
|  | qSW6 | 6 | S6_5982269 | S6_41973878 | 0 | 6.89 | 5 | 6.89 | 4.86 | 5.31 | 0.06 |
|  | qSW7 | 7 | S7_55367570 | S7_56145255 | 60.43 | 62.13 | 61 | 1.70 | 9.32 | 10.89 | -0.09 |

LOD, log<sub>10</sub> of the likelihood odds ratio; PVE (%), phenotypic variation explained by the marker; Add, the additive effect of QTL across four environments.

**Fig. 2.**
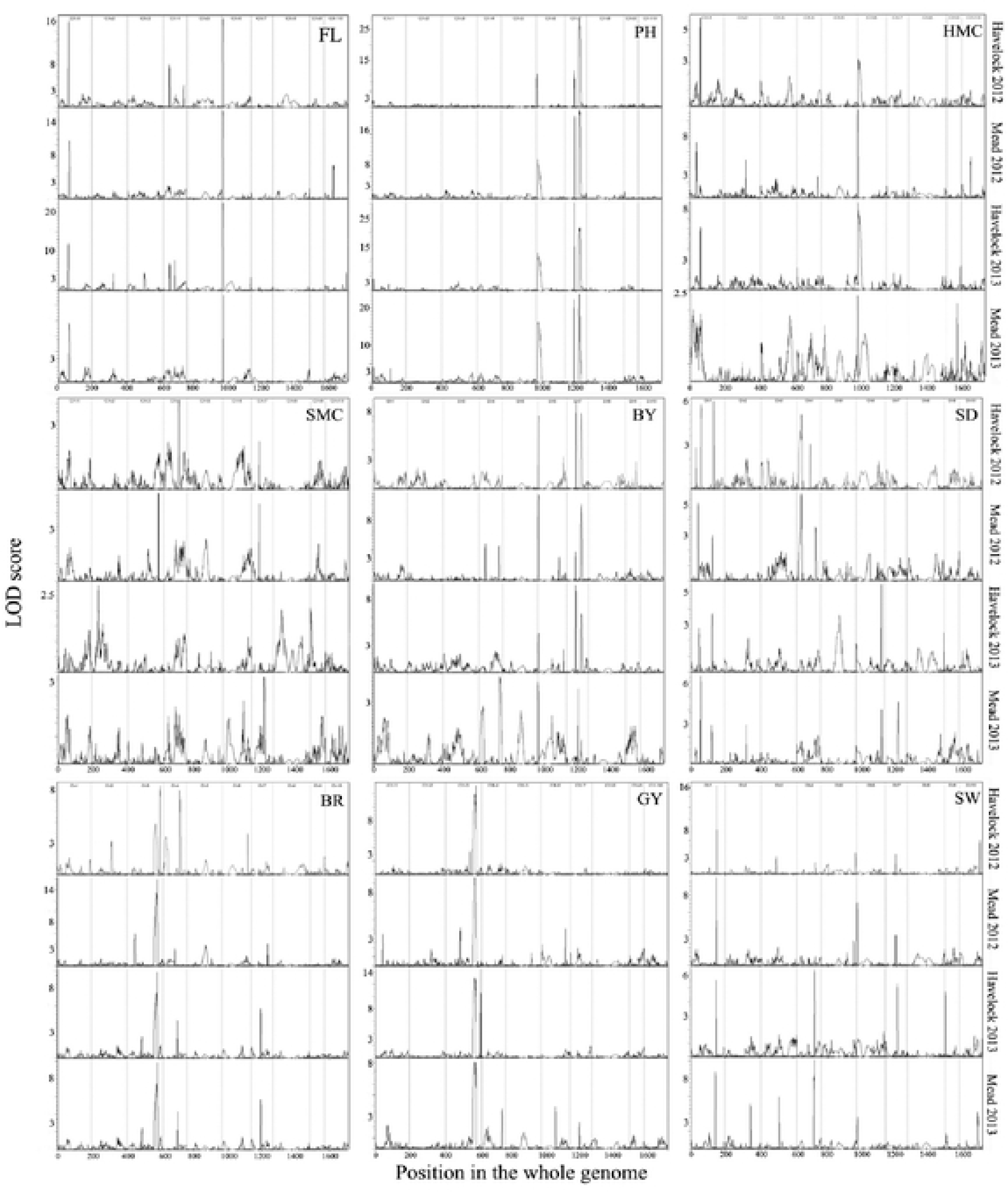
QTL locations in the linkage map for nine bioenergy-related traits across four environments. FL, flowering time; PH, plant height; HMC, head moisture content; SMC, stem moisture content; BY, total biomass yield; SD, stem diameter; BR, brix; GY, grain yield; SW, 100 seed weight; LOO, log_10_ of the likelihood odds r

**Fig. 3.**
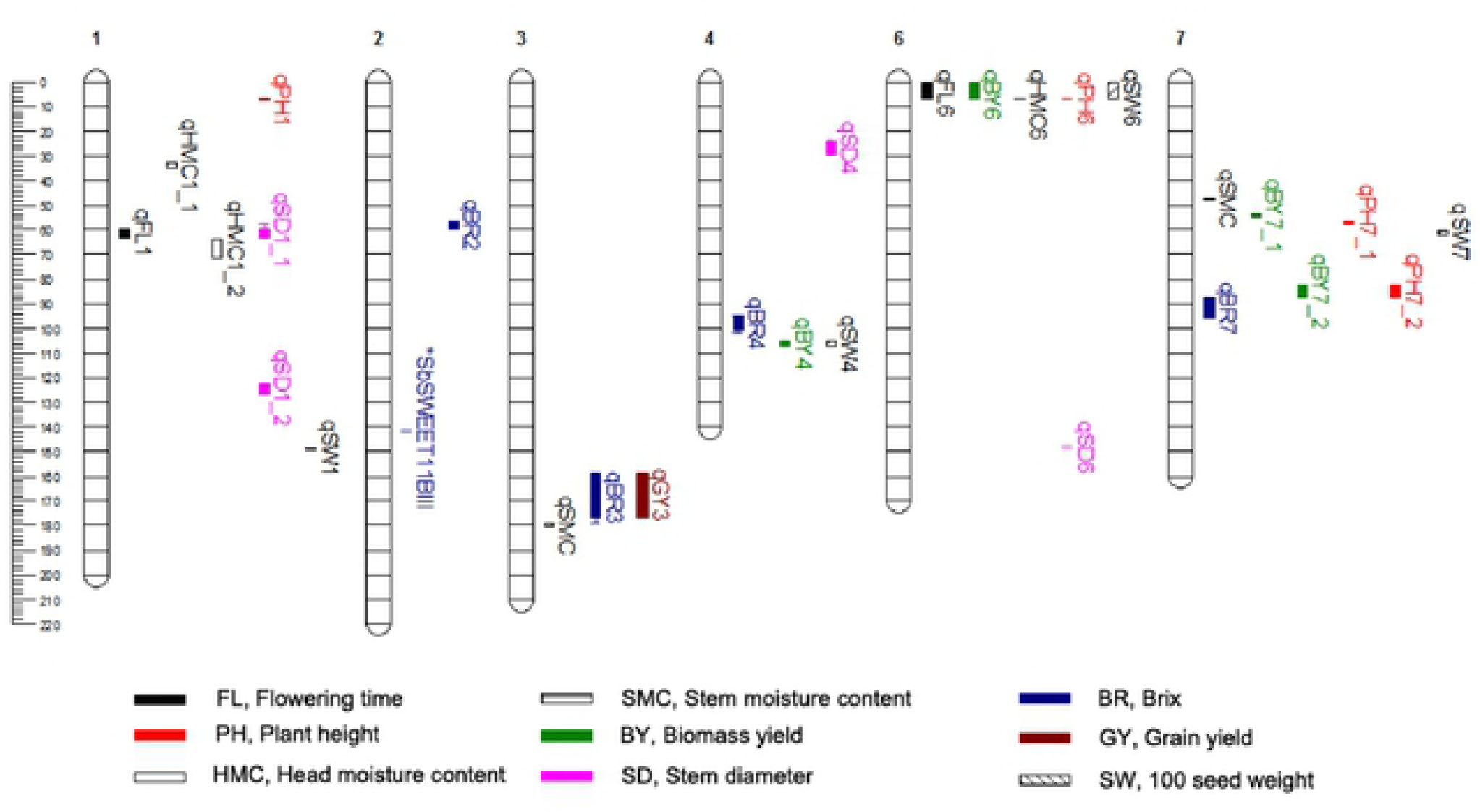
QTL locations in the linkage map for nine bioenergy-related traits across four environments

### QTLs for flowering time

A major QTL is defined as a QTL that explains >10% PVE. There were two major QTLs associated with flowering time, located on chromosomes 1 and 6. The major QTL on chromosome 1 explained 18.17% PVE and had a positive additive effect (3.63 days). The major QTL with the highest PVE for flowering time was on chromosome 6 (29.45% PVE) with a negative additive effect (−4.66 days) and was detected consistently across three environments (Havelock 2012, Mead 2012 and Havelock 2013).

### QTLs for plant height

A total of four QTLs were detected for plant height on chromosomes 1, 6, and 7, and explained 3.10-29.51% PVE. Of the four QTLs, three QTLs on chromosome 6 and 7 were major QTLs and had negative effects ranging from −17.53 to −21.68 cm, while the QTL on chromosome 1 had an 8.04 cm additive effect. The major QTL with the highest PVE (29.51%) was on chromosome 7 and was detected consistently across four environments.

### QTLs for head moisture content

There were three QTLs associated with head moisture content, which were located on chromosomes 1 and 6, and explained 7.15-20.65% PVE. Their additive effects were 1.81% and 1.73% for two QTLs on chromosome 1, and −2.92% for a QTL on chromosome 6. The major QTL with the highest PVE (20.65%) was on chromosome 6, but was detected only at Mead in 2012 and Havelock in 2013. None of the QTLs associated with head moisture were detected at Mead in 2013.

### QTLs for stem moisture content

None of the QTLs for stem moisture content were detected for combined environments, but two major QTLs for stem moisture content were detected individually at Mead in 2012 on chromosomes 3 and 7. The QTL located on chromosome 3 explained 10.77% PVE and had a 0.75% additive effect, while the QTL located on chromosome 7 explained 10.29% PVE and had a −0.74% additive effect (Appendix 2).

### QTLs for biomass yield

Four QTLs were detected for total biomass yield on chromosomes 4, 6, and 7, and explained 5.72-16.46% PVE. Of the four QTLs, three QTLs on chromosomes 6 and 7 were major QTLs. The major QTL on chromosome 7 (flanking markers; S7_58245093-58601476) was also the major QTL identified for plant height. The QTL with the highest % PVE (16.46%) was on chromosome 7 and was detected consistently across three environments (Havelock 2012, Mead 2012, and Havelock 2013). There was a major QTL located on chromosome 6 (10.93% PVE) that was detected consistently across four environments. All QTLs for total biomass yield had negative additive effects ranging from −0.97 to −1.64 Mg ha^-1^.

### QTLs for stem diameter

Five QTLs for stem diameter were detected on chromosomes 1, 4, and 6, and explained 4.81-9.43 % PVE. The QTL with the highest PVE was on chromosome 1 but was detected only at Havelock in 2012. QTLs on chromosomes 1 and 4 had positive additive effects with 0.05 cm for all QTLs, while a QTL on chromosome 6 had a negative additive effect (−0.04 cm).

### QTLs for brix

There were six QTLs associated with brix, which were located on chromosomes 2, 3, 4, and 7 and explained 5.28-39.92% PVE. Two QTLs on chromosome 3 were major QTLs. The major QTL with the highest PVE was on chromosome 3 (39.92% PVE) and was detected consistently across all four environments. There were negative additive effects of the QTLs detected on chromosomes 2, 3, and 7 and ranged from −0.62 to −1.71. QTLs detected on chromosome 4 had positive additive effects ranging from 0.79-0.82.

### QTL for grain yield

Only one major QTL for grain yield was detected on chromosome 3 with 49.14% PVE and was detected consistently across four environments. This major QTL co-localized with the highest PVE QTL for brix, and its additive effect was 0.97 Mg ha^-1^.

### QTLs for 100 seed weight

There were four QTLs associated with 100 seed weight, which were located on chromosomes 1, 4, 6, and 7, and explained 5.31-22.97% PVE. QTLs on chromosome 1 and 7 were major QTLs. The major QTL with the highest PVE was on chromosome 1 (22.97% PVE) but was detected only at Havelock in 2013. There were positive additive effects of the QTLs detected on chromosomes 1, 4, and 6 and ranged from 0.06-0.13 g, while the QTL detected on chromosome 7 had a negative additive effect (−0.09 g).

## Discussion

### The sorghum genetic map

The TASSEL4 GBS pipeline provided the HapMap file containing 10,424 SNPs but only 979 SNPs were polymorphic SNPs used for linkage map construction. The results were similar to Kong et al. (2018) and Gelli et al. (2016) that identified GBS based polymorphic SNP markers in sorghum RIL for linkage map construction with 616 and 642 polymorphic SNPs and the linkage maps spanned 1404.8 and 1641 cM., respectively. The linkage map for this study spanned 1,707.11 cM, which is comparable with previously reported sorghum genetic maps that varied from 603.5 to 2,128 (Gelli et al. 2016; Jordan et al. 2011; Kong et al. 2018). The average inter-marker distance was 1.74 cM, showing a high quality genetic map using GBS in RIL population.

### QTLs for flowering time associated with *Ma1* and *SbEHD1*

Sorghum flowering time or maturity genes were identified at six loci, designated *Ma1* on chromosome 6 (locus name: *SbPRR37*, Sobic.006G057866; region: 40304883-40316799), *Ma2* on chromosome 2 (unknown), *Ma3* on chromosome 1 (*SbPHYB*, Sobic.001G394400: 68034103-68043358), *Ma4* on chromosome 10 (unknown), *Ma5* on chromosome 1 (*SbPHYC*, Sobic.001G087100: 6748036-6753421), and *Ma6* on chromosome 6 (*SbGHD7*, Sobic.006G004400: 697459-700101) (Childs et al. 1997; Mullet and Rooney 2013; Murphy et al. 2014; Murphy et al. 2011; Quinby 1966; Rooney and Aydin 1999; White et al. 2004; Yang et al. 2014). We detected a major QTL for flowering time on chromosome 6 (QTL name in this study: AD6; flanking markers: S6_5982269-41974107; position: 0-6.89 cM) which contains *Ma1*. Using the Phyzotome database (www.phytozome.net) to browse annotated genes for *Sorghum bicolor* v3.1, we found the closest SNP (S6_41974107) located 1.66 Mbp from the right side of *Ma1* (40304883-40316799). Recently, Yang et al. (2014) reported that *SbEHD1*, an activator of flowering in sorghum, is located on chromosome 1 near this region (Sobic.001G227900: 21860030-21867056). In this study, a major QTL for flowering time was detected on chromosome 1 (AD1, S1_19760428-20278787, 59.63-62.88 cM) and the closest SNP (S1_20278787) was located 1.58 Mbp to the left of *SbEHD1* (21860030-2186705). These results suggest that QTLs for flowering time on chromosomes 6 and 1 may be contributed by *Ma1* and *SbEHD1*, respectively.

Similar results were also observed in previous QTL mapping for flowering time in sweet x grain sorghum populations. Ritter et al. (2008) reported a QTL for flowering time on chromosome 1 associated with marker ACC/CA3 or at 40.1 cM, whereas the peak position of the QTL located on chromosome 1 in this study was at 61 cM. In previous studies, QTLs for flowering time in sweet sorghum were mostly mapped on chromosome 6 at 0-48 cM, which contains the *Ma1 and Ma6* genes (Felderhoff et al. 2012; Murray et al. 2008b; Ritter et al. 2008; Shiringani et al. 2010).

### QTLs for plant height associated with *dw2* and *dw3*

Plant height in sorghum is influenced by four loci, *dw1* on chromosome 9 (Sobic.009G229800: 57038653-57041166), *dw2* on chromosome 6 (unknown), *dw3* on chromosome 7 (Sobic.007G163800: 59821905-59829910), and *dw4* on chromosome 6 (unknown) (Multani et al. 2003; Quinby 1974; Yamaguchi et al. 2016). Only *dw1* and *dw3* have been cloned and sequenced. A major QTL for plant height in the current study was identified on chromosome 7 (PH7_2, S7_58245093-58601476, 82.13-87.37 cM). The closest SNP (S7_58601476) was located 1.22 Mbp to the left of *dw3*. QTLs for plant height in this study were also identified on chromosomes 1 and 6. Similarly, QTLs for plant height in sweet sorghum have been reported from previous studies on chromosome 1 (Guan et al. 2011; Liu et al. 2015; Ritter et al. 2008; Shiringani et al. 2010), chromosome 6 (Felderhoff et al. 2012; Murray et al. 2008b; Ritter et al. 2008; Shiringani et al. 2010), and chromosome 7 (Felderhoff et al. 2012; Guan et al. 2011; Murray et al. 2008b; Shehzad and Okuno 2015; Shiringani et al. 2010). The QTLs for plant height reported from previous studies and this study were located in nearby regions on chromosomes 1 (Liu et al. 2015), 6 (Ritter et al. 2008; Shiringani et al. 2010), and 7 (Felderhoff et al. 2012; Murray et al. 2008b; Shiringani et al. 2010). The QTL on chromosome 6 (PH6, S6_41974107-42311647, 6.89-8.56 cM) located at 41.97-42.31 Mb co-localizes with the QTL for temperate plant height (41-43 Mb) identified by Higgins *et al*. (2014). This region is also the putative region containing the gene *dw2* reported by recent studies using GBS (Gelli *et al*., 2016, 41.97-43.22 Mb; Higgins *et al*., 2014, 41.23-44.34 Mb; Morris *et al*., 2013, 42.26 Mb). These results indicated that QTLs for plant height on chromosomes 6 and 7 in this study may be contributed by *dw2* and *dw3*, respectively. The QTLs on chromosomes 6 and 7 were major QTLs and had negative additive effects showing that *dw2* and *dw3* were transferred from Macia, the short grain sorghum parent line used in this study.

### Co-localization of QTLs for flowering time, plant height, and biomass yield

Plant height and flowering time are highly related traits in grasses, in which apical growth is terminated by flowering (Lin et al. 1995). QTLs associated with plant height are linked with loci controlling flowering, and have been previously reported in wheat (Börner et al. 1993), maize (Khairallah et al. 1998; Thornsberry et al. 2001), rice (Yan et al. 2011), barley (Chen et al. 2009), and sorghum (Börner et al. 1993; Higgins et al. 2014; Khairallah et al. 1998; Lin et al. 1995; Madhusudhana and Patil 2013; Morris et al. 2013; Shiringani et al. 2010; Thornsberry et al. 2001). In this study, a QTL for plant height (PH6, S6_41974107-42311647, 6.89-8.56 cM) co-localized with a major QTL for flowering time on chromosome 6. This result supports the finding that the dwarfing locus, *dw2*, is linked to the maturity gene, *Ma1*, which had been investigated in sorghum (Higgins et al. 2014; Lin et al. 1995; Madhusudhana and Patil 2013; Morris et al. 2013; Shiringani et al. 2010). We also found that total biomass yield correlated best with plant height (*r*=0.67), and major QTLs for these traits co-localized on chromosomes 6 and 7. Similarly, Ritter et al. (2008), Shiringani et al. (2010), and Shiringani and Friedt (2011) also reported the co-localization on chromosome 6, while Murray et al. (2008b) and Felderhoff et al. (2012) reported the co-localization on chromosome 7. The results indicated that total biomass yield is highly determined by plant height.

### QTLs for brix associated with *SbSWEET1A, SbSUT5*, and *SbSUT6*

Sugar content in this study was measured using brix. Brix is used as a practical analysis of total solute content (mostly sugars) (Kawahigashi et al. 2013). Ritter et al. (2008) suggested that brix is a simpler phenotypic trait compared with quantifying sucrose and sugar content, which are more time-consuming and expensive to measure. Moreover, they identify similar QTLs. In the current study, QTLs for brix were identified on chromosomes 2, 3, 4, and 7. Similar results were also observed in previous QTL mapping for brix on chromosome 2 (Guan et al. 2011; Shiringani et al. 2010), chromosome 3 (Felderhoff et al. 2012; Guan et al. 2011; Murray et al. 2008b), chromosome 4 (Bian et al. 2006; Felderhoff et al. 2012; Lekgari 2010; Shiringani et al. 2010), and chromosome 7 (Bian et al. 2006; Guan et al. 2011; Lekgari 2010; Murray et al. 2008b; Shiringani et al. 2010). The QTLs for brix identified in the present study may be associated with known candidate genes controlling sugar content in sorghum. Genes suggested to control the accumulation of sugar in sorghum stems have been characterized, including six sucrose transporter genes (*SbSUTs*), three tonoplast sugar transporter genes (*SbTSTs*) and twenty SWEET genes (*SbSWEETs*), (Bihmidine et al. 2015; Bihmidine et al. 2016; Braun and Slewinski 2009; Eom et al. 2015; Milne et al. 2013; Qazi et al. 2012). Intriguingly, we found that the major QTL detected on chromosome 3 (BR3_1, S3_68877230-69154987, 158.76-176.55 cM) was located only 4.1 kbp from the left side of *SbSWEET1A* (Sobic.003G377700: 69196094-69199768), suggesting that *SbSWEET1A* could be the candidate gene underlying the QTL. The QTL we detected on chromosome 4 (BR4_1, S4_56025428-57727357, 94.27-100.68 cM) was located 1.80 Mbp from the right side of *SbSUT5* (Sobic.004G190500: 54229192-54232507), and was also potentially identified by Shiringani et al. (2010). The QTL we detected on chromosome 7 (BR7, S7_58601476-60256664, 87.37-95.91 cM) was located 4.04 Mbp from the left side of *SbSUT6* (Sobic.007G214500: 64300794-64303945). A similar QTL was reported by Murray et al. (2009) and Lekgari (2010). These results suggest that the genes *SbSWEET1A, SbSUT5*, and *SbSUT6* may underlie the QTLs for brix identified on chromosomes 3, 4, and 7. Interestingly, Milne et al. (2013) reported that *SbSUT5* and *SbSUT6* may function in sucrose phloem loading in leaves and sucrose accumulation in stems. However, a previous qRT-PCR expression analysis of *SbSUT5* and *SbSUT6* in Wray and Macia stems did not support the hypothesis that these genes play a prominent role in sucrose phloem loading or stem sugar accumulation (Bihmidine et al. 2015). Further functional analyses of these genes will determine their contributions to whole-plant carbohydrate partitioning and sugar accumulation in stems.

It is also noteworthy, although we did not detect similar QTLs in our analyses, several of the other studies identified QTLs for brix that overlap the position for sugar transporter genes. For example, on chromosome 2, Guan et al. (2011) found a QTL for brix near the *SbSWEET11B* gene, and on chromosome 3, Felderhoff et al. (2012) and Guan et al. (2011) identified QTL containing the *SbSWEET2A* and *SbSWEET6* genes. Additionally, on chromosome 4, Bian et al. (2006), Lekgari (2010), Shiringani et al. (2010) and Felderhoff et al. (2012) identified brix QTL that are in the vicinity of *SbSWEET15, SbSWEET4A, SbSWEET4B, SbSWEET4C*, and *SbTST2*. Future research will investigate the molecular functions of these genes and their potential involvement in regulating stem sugar content.

Finally, we also found another QTL for brix on chromosome 2 that was not correlated with known sucrose transport genes in sorghum. Shiringani et al. (2010) reported a QTL for brix on chromosome 2 at positions 30-40 cM, while the QTL for brix located on chromosome 2 in this study was at position 56.3-59.4 cM. It is possible that these QTLs for brix will be contributors for identifying other sugar transporter genes or novel genes affecting sugar metabolism.

### Pleiotropic effects of QTLs associated with brix and grain yield

A major QTL for brix and grain yield was co-localized on chromosome 3 (BR3_1 and GY3, S3_62297204-68877230, 158.76-176.55 cM. Felderhoff et al. (2012) also reported QTLs for brix and grain yield located nearby genomic regions on chromosome 3. Pleiotropic effects of genes that control more than one quantitative trait might explain this result. On the other hand, two or more nearby genes that control different traits might explain the results of co-localized regions. It is interesting to note that a related *SWEET* gene in maize and rice controls grain filling (Sosso et al. 2015). In our study, brix was negatively correlated with grain yield (*r*=-0.39) as expected for competing sinks (Bihmidine et al. 2013). Sweet sorghum varieties that traditionally have high concentration of sugar in stems have a small panicle because of competition for carbohydrates between grain filling and sugar storage in stems (Bihmidine et al. 2015; Gutjahr et al. 2013).

### QTLs for stem moisture content

QTLs for all traits were detected for combined environments, except QTLs for stem moisture content. The reliability of the confidence intervals associated with QTL locations depends on the heritability of the individual QTL (Kearsey and Farquhar 1998). QTLs related to stem moisture content could not be detected in combined environments, but they were detected in one environment (Mead 2012) on chromosomes 3 and 7. Stem moisture content is easily affected by environment. The heritability of stem moisture content for combined environments was as low as 0.25, whereas that of Mead in 2012 was 0.48. The low heritability of the stem moisture content QTLs is likely the reason that we failed to detect them in the analysis. This result is similar to that found by Felderhoff et al. (2012). They reported no QTLs for percent moisture across all environments, but they found a QTL for moisture on chromosome 3 at only one of four locations.

QTLs detected in this study were similar in genomic regions with previous studies. Some QTLs were located on the same position, and genomic regions were narrowed in this study. QTLs for head moisture content were identified on chromosome 1, which is similar to the data of Lekgari (2010), who reported a QTL related to head moisture content on chromosome 1 at position 52.6 cM. In this study, QTLs for this trait were detected on chromosome 1 at positions 34 and 65 cM. QTLs for stem diameter were detected on chromosomes 1, 4, and 6. Two QTLs on chromosome 1 were located at the same position as QTLs reported by Shehzad and Okuno (2015) and by Shiringani et al. (2010), but both regions from this study were smaller than the previous studies.

Similarly, for 100 seed weight, we detected QTLs on the same chromosomes as found in previous studies, but ours were located at different positions. In this study, QTLs for 100 seed weight were detected on chromosomes 1, 4, 6, and 7 at positions 148.7-149.3, 105.1-107.4, 0-6.9, and 60.4-62.1 cM, respectively. Han et al. (2015) and Murray et al. (2008b) found QTLs for 100 seed weight at overlapping regions of 0-19 cM on chromosome 1. Shehzad and Okuno (2015) reported two QTLs for 100-grain weight on chromosome 4 at position 0-12 cM and 17.9-30.9 cM. Murray et al. (2008b) found a QTL for 1000 seed weight on chromosome 6 at position 9.6-30.8 cM. Han et al. (2015) and Shehzad and Okuno (2015) also reported QTLs for 100 seed weight on chromosome 7 at position 150.9-161.4 and 76.3-93.6 cM, respectively. These differences are likely attributable to differences in genotypes evaluated and environmental conditions.

In conclusion, GBS increased the precision of the QTL analysis and related genes could be identified. The bioenergy-related QTLs in sweet sorghum detected in this study were located in genomic regions associated with known genes linked to the traits (flowering time, plant height, and brix). Other QTLs we identified overlap with previous studies. Tantalizingly, the association between *SbSWEET1A* and the major QTL for brix on chromosome 3 suggests a potential mechanism underlying sugar accumulation in the sorghum (and possibly other bioenergy grasses) stem. Future validation of the bioenergy-related QTLs and identification of associated candidate genes should be conducted for functional genomic improvement of bioenergy-related traits in sweet sorghum.

## Abbreviations

FL: Flowering time
PH: Plant height
HMC: Head moisture content
SMC: Stem moisture content
BY: Biomass yield
SD: Stem diameter
BR: Brix
GY: Grain yield
SW: 100 Seed weight

## Acknowledgments

We are grateful to Benjamin Babst, Brookhaven national laboratory for collaboration. This work is being funded by a grant from the Plant Feedstock Genomics for Bioenergy # DE-SC0006810 to DMB et al.

## References

Beissinger TM et al. (2013) Marker density and read depth for genotyping populations using genotyping-by-sequencing Genetics 193:1073–1081

Bian Yl, Yazaki S, Inoue M, Cai Hw (2006) QTLs for sugar content of stalk in sweet sorghum (*Sorghum bicolor* L. Moench) Agricultural Sciences in China 5:736–744

Bihmidine S, Baker RF, Hoffner C, Braun DM (2015) Sucrose accumulation in sweet sorghum stems occurs by apoplasmic phloem unloading and does not involve differential Sucrose transporter expression BMC plant biology 15:186

Bihmidine S, Hunter III CT, Johns CE, Koch KE, Braun DM (2013) Regulation of assimilate import into sink organs: update on molecular drivers of sink strength Frontiers in plant science 4:177

Bihmidine S, Julius B, Dweikat I, Braun DM (2016) Tonoplast Sugar Transporters (SbTSTs) putatively control sucrose accumulation in sweet sorghum stems Plant signaling & behavior:00-00

Börner A, Worland A, Plaschke J, Schumann E, Law C (1993) Pleiotropic effects of genes for reduced height (Rht) and day-length insensitivity (Ppd) on yield and its components for wheat grown in middle Europe Plant Breeding 111:204–216

Bradbury PJ, Zhang Z, Kroon DE, Casstevens TM, Ramdoss Y, Buckler ES (2007) TASSEL: Software for association mapping of complex traits in diverse samples Bioinformatics 23:2633–2635

Braun DM, Slewinski TL (2009) Genetic control of carbon partitioning in grasses: roles of sucrose transporters and tie-dyed loci in phloem loading Plant Physiology 149:71–81

Broadhead D, Freeman K, Zummo N (1978) ‘Wray’-a new variety of sweet sorghum for sugar production [Yields] Research Report-Mississippi Agricultural and Forestry Experiment Station (USA) no 4 (1)

Broman KW, Sen S (2009) A Guide to QTL Mapping with R/qtl vol 46. Springer,

Calviño M, Messing J (2012) Sweet sorghum as a model system for bioenergy crops Current Opinion in Biotechnology 23:323–329 DOI:http://dx.doi.org/10.1016/j.copbio.2011.12.002

Carpita NC, McCann MC (2008) Maize and sorghum: genetic resources for bioenergy grasses Trends in Plant Science 13:415–420 DOI:http://dx.doi.org/10.1016/j.tplants.2008.06.002

Chen A, Baumann U, Fincher GB, Collins NC (2009) Flt-2L, a locus in barley controlling flowering time, spike density, and plant height Functional & integrative genomics 9:243–254

Childs KL, Miller FR, Cordonnier-Pratt M-M, Pratt LH, Morgan PW, Mullet JE (1997) The sorghum photoperiod sensitivity gene, Ma3, encodes a phytochrome B Plant Physiology 113:611–619

Demirbas A (2001) Biomass resource facilities and biomass conversion processing for fuels and chemicals Energy Conversion and Management 42:1357–1378

Elshire RJ, Glaubitz JC, Sun Q, Poland JA, Kawamoto K, Buckler ES, Mitchell SE (2011) A robust, simple genotyping-by-sequencing (GBS) approach for high diversity species PLoS ONE 6:e19379 DOI:10.1371/journal.pone.0019379

Eom J-S et al. (2015) SWEETs, transporters for intracellular and intercellular sugar translocation Current opinion in plant biology 25:53–62

Felderhoff TJ et al. (2012) QTLs for energy-related traits in a sweet × grain sorghum [*Sorghum bicolor* (L.) Moench] mapping population Crop Science 52:2040–2049

Gelli M et al. (2016) Mapping QTLs and association of differentially expressed gene transcripts for multiple agronomic traits under different nitrogen levels in sorghum BMC plant biology 16:1

Guan Y et al. (2011) QTL mapping of bio-energy related traits in sorghum Euphytica 182:431–440

Guo M, Song W, Buhain J (2015) Bioenergy and biofuels: History, status, and perspective Renewable and Sustainable Energy Reviews 42:712–725

Gutjahr S, Vaksmann M, Dingkuhn M, Thera K, Trouche G, Braconnier S, Luquet D (2013) Grain, sugar and biomass accumulation in tropical sorghums. I. Trade-offs and effects of phenological plasticity Functional Plant Biology 40:342–354 DOI:10.1071/FP12269

Han L et al. (2015) Fine mapping of qGW1, a major QTL for grain weight in sorghum Theoretical and Applied Genetics 128:1813–1825

Higgins RH, Thurber CS, Assaranurak I, Brown PJ (2014) Multiparental mapping of plant height and flowering time QTL in partially isogenic sorghum families G3: Genes, Genomes, Genetics 4:1593–1602 DOI:10.1534/g3.114.013318

Hills F, Lewellen R, Skoyen I (1990) Sweet sorghum cuItivars for alcohol production California Agriculture 44:14–16

Holland JB, Nyquist WE, Cervantes-Martínez CT (2003) Estimating and interpreting heritability for plant breeding: An update Plant Breeding Reviews 22:9–112

Jordan DR, Mace ES, Cruickshank AW, Hunt CH, Henzell RG (2011) Exploring and exploiting genetic variation from unadapted sorghum germplasm in a breeding program Crop Sci 51:1444–1457 DOI:10.2135/cropsci2010.06.0326

Kawahigashi H, Kasuga S, Okuizumi H, Hiradate S, Yonemaru J-i (2013) Evaluation of Brix and sugar content in stem juice from sorghum varieties Grassland Science 59:11–19 DOI:10.1111/grs.12006

Kearsey M, Farquhar A (1998) QTL analysis in plants; where are we now? Heredity 80:137–142

Khairallah M et al. (1998) Molecular mapping of QTL for southwestern corn borer resistance, plant height and flowering in tropical maize Plant Breeding 117:309–318

Kong W et al. (2018) Genotyping by Sequencing of 393 *Sorghum bicolor* BTx623 × IS3620C recombinant inbred lines improves sensitivity and resolution of QTL Detection G3: Genes|GenomesGenetics 8:2563–2572 DOI:10.1534/g3.118.200173

Lekgari AL (2010) Genetic mapping of quantitative trait loci associated with bioenergy traits, and the assessment of genetic variability in sweet sorghum (Sorghum bicolor (L.). Moench)

Li H, Durbin R (2009) Fast and accurate short read alignment with Burrows-Wheeler transform Bioinformatics 25:1754–1760 DOI:10.1093/bioinformatics/btp324

Lin YR, Schertz KF, Paterson AH (1995) Comparative analysis of QTLs affecting plant height and maturity across the poaceae, in reference to an interspecific sorghum population Genetics 141:391–411

Littell RC, Stroup WW, Milliken GA, Wolfinger RD, Schabenberger O (2006) SAS for mixed models. SAS institute,

Liu YL, Wang LH, Li JQ, Zhan QW, Zhang Q, Li JF, Fan FF (2015) QTL mapping of forage yield and forage yield component traits in Sorghum bicolor x S. sudanense Genetics and Molecular Research 14:3854–3861 DOI:10.4238/2015.April.22.14

Lv P et al. (2013) Association analysis of sugar yield-related traits in sorghum *Sorghum bicolor* (L.) Euphytica 193:419–431 DOI:10.1007/s10681-013-0962-7

Madhusudhana R (2014) Genetic mapping in sorghum genetics, Genomics and Breeding of Sorghum:141

Madhusudhana R, Patil JV (2013) A major QTL for plant height is linked with bloom locus in sorghum [*Sorghum bicolor* (L.) Moench] Euphytica 191:259–268

Makanda I, Tongoona P, Derera J (2009) Combining ability and heterosis of sorghum germplasm for stem sugar traits under off-season conditions in tropical lowland environments Field Crops Research 114:272–279 DOI:http://dx.doi.org/10.1016/j.fcr.2009.08.009

Milne RJ, Byrt CS, Patrick JW, Grof CPL (2013) Are sucrose transporter expression profiles linked with patterns of biomass partitioning in Sorghum phenotypes? Frontiers in Plant Science 4:223 DOI:10.3389/fpls.2013.00223

Morris GP et al. (2013) Population genomic and genome-wide association studies of agroclimatic traits in sorghum Proceedings of the National Academy of Sciences of the United States of America 110:453–458 DOI:10.1073/pnas.1215985110

Mullet JE, Rooney WL (2013) Method for production of sorghum hybrids with selected flowering times. Google Patents,

Multani DS, Briggs SP, Chamberlin MA, Blakeslee JJ, Murphy AS, Johal GS (2003) Loss of an MDR transporter in compact stalks of maize br2 and sorghum dw3 mutants Science 302:81–84 DOI:10.1126/science.1086072

Murphy R, Morishige D, Brady J, Rooney W, Yang S, Klein P, Mullet J (2014) Ghd7 (Ma6) Represses flowering in long days: a key trait in energy sorghum hybrids Plant Genome

Murphy RL et al. (2011) Coincident light and clock regulation of pseudoresponse regulator protein 37 (PRR37) controls photoperiodic flowering in sorghum Proceedings of the National Academy of Sciences 108:16469–16474

Murray SC, Rooney WL, Hamblin MT, Mitchell SE, Kresovich S (2009) Sweet sorghum genetic diversity and association mapping for brix and height plant genome 2:48–62 DOI:10.3835/plantgenome2008.10.0011

Murray SC, Rooney WL, Mitchell SE, Sharma A, Klein PE, Mullet JE, Kresovich S (2008a) Genetic improvement of sorghum as a biofuel feedstock: II. QTL for stem and leaf structural carbohydrates Crop Sci 48:2180–2193 DOI:10.2135/cropsci2008.01.0068

Murray SC, Sharma A, Rooney WL, Klein PE, Mullet JE, Mitchell SE, Kresovich S (2008b) Genetic improvement of sorghum as a biofuel feedstock: I. QTL for stem sugar and grain nonstructural carbohydrates Crop Sci 48:2165–2179 DOI:10.2135/cropsci2008.01.0016

Paterson AH et al. (2009) The *Sorghum bicolor* genome and the diversification of grasses Nature 457:551–556 DOI:10.1038/nature07723

Pedersen JF, Sattler SE, Anderson WF (2013) Evaluation of public sweet sorghum A-lines for use in hybrid production Bioenerg Res 6:91–102

Qazi HA, Paranjpe S, Bhargava S (2012) Stem sugar accumulation in sweet sorghum–activity and expression of sucrose metabolizing enzymes and sucrose transporters Journal of plant physiology 169:605–613

Quinby JR (1966) Fourth maturity gene locus in sorghum Crop Sci 6:516–518

Quinby JR (1974) Sorghum improvement and the genetics of growth

Ritter KB, Jordan DR, Chapman SC, Godwin ID, Mace ES, Lynne McIntyre C (2008) Identification of QTL for sugar-related traits in a sweet x grain sorghum (*Sorghum bicolor* L. Moench) recombinant inbred population Molecular Breeding 22:367–384

Rooney WL, Aydin S (1999) Genetic control of a photoperiod-sensitive response in *Sorghum bicolor* (L.) Moench Crop Science 39:397–400

Rooney WL, Blumenthal J, Bean B, Mullet JE (2007) Designing sorghum as a dedicated bioenergy feedstock Biofuels, Bioproducts and Biorefining 1:147–157

Saadan H, Mgonja M, Obilana A (2000) Performance of the sorghum variety Macia in multiple environments in Tanzania International Sorghum and Millets Newsletter 41:10–12

Setimela P, Manthe C, Mazhani L, Obilana A (1997) Release of three grain sorghum pure line varieties in Botswana South African Journal of Plant and Soil 14:137–138

Shehzad T, Okuno K (2014) Diversity assessment of sorghum germplasm and its utilization in genetic analysis of quantitative traits-A review Australian Journal of Crop Science 8:937–944

Shehzad T, Okuno K (2015) QTL mapping for yield and yield-contributing traits in sorghum (*Sorghum bicolor* (L.) Moench) with genome-based SSR markers Euphytica 203:17–31 DOI:10.1007/s10681-014-1243-9

Shiringani AL, Friedt W (2011) QTL for fibre-related traits in grain × sweet sorghum as a tool for the enhancement of sorghum as a biomass crop Theoretical and Applied Genetics 123:999–1011

Shiringani AL, Frisch M, Friedt W (2010) Genetic mapping of QTLs for sugar-related traits in a RIL population of *Sorghum bicolor* L. Moench Theoretical and Applied Genetics 121:323–336

Sosso D et al. (2015) Seed filling in domesticated maize and rice depends on SWEET-mediated hexose transport Nature genetics

Thornsberry JM, Goodman MM, Doebley J, Kresovich S, Nielsen D, Buckler ES (2001) Dwarf8 polymorphisms associate with variation in flowering time Nature genetics 28:286–289

Verma S, Gupta S, Bandhiwal N, Kumar T, Bharadwaj C, Bhatia S (2015) High-density linkage map construction and mapping of seed trait QTLs in chickpea (Cicer arietinum L.) using Genotyping-by-Sequencing (GBS) Scientific Reports 5:17512 DOI:10.1038/srep17512 https://www.nature.com/articles/srep17512#supplementary-information

Wang J, Li H, Zhang L, Meng L (2012) QTL IciMapping version 3.2 The Quantitative Genetics Group Institute of Crop Science Chinese Academy of Agricultural Sciences (CAAS) Beijing 100081

White GM, Hamblin MT, Kresovich S (2004) Molecular evolution of the phytochrome gene family in sorghum: changing rates of synonymous and replacement evolution Molecular biology and evolution 21:716–723

Xin Z, Chen J (2012) A high throughput DNA extraction method with high yield and quality Plant Methods 8

Yamaguchi M et al. (2016) Sorghum Dw1, an agronomically important gene for lodging resistance, encodes a novel protein involved in cell proliferation Scientific Reports 6:28366 DOI:10.1038/srep28366

Yan W-H et al. (2011) A major QTL, Ghd8, plays pleiotropic roles in regulating grain productivity, plant height, and heading date in rice Molecular Plant 4:319–330

Yang S, Murphy RL, Morishige DT, Klein PE, Rooney WL, Mullet JE (2014) Sorghum Phytochrome B Inhibits Flowering in Long Days by Activating Expression of SbPRR37 and SbGHD7, Repressors of SbEHD1, SbCN8 and SbCN12 Plos One 9 DOI:10.1371/journal.pone.0105352

Zegada-Lizarazu W, Monti A (2012) Are we ready to cultivate sweet sorghum as a bioenergy feedstock? A review on field management practices Biomass and Bioenergy 40:1–12 DOI: http://dx.doi.org/10.1016/j.biombioe.2012.01.048

